# Spatiotemporal changes in genetic diversity and structure of a recent fish invasion in eastern North America

**DOI:** 10.1101/2022.03.01.482476

**Authors:** Thaïs A. Bernos, Sunčica Avlijaš, Jaclyn Hill, Olivier Morissette, Anthony Ricciardi, Nicholas E. Mandrak, Kenneth M. Jeffries

## Abstract

Introduced and geographically expanding populations experience similar eco-evolutionary challenges, including founder events, genetic bottlenecks, and novel environments. Theory predicts that reduced genetic diversity resulting from such genetic phenomena limits the colonization success of introduced populations. We examined an invasive population of a Eurasian freshwater fish, Tench (*Tinca tinca*), that has been expanding geographically in eastern North America for three decades. Using genomic data, we evaluated evidence for single versus multiple introductions and the connectivity of the population across the entire range in which it has been spreading. Tench exhibited low levels of genetic diversity, a lack of marked population subdivision across time and space, and evidence of a recent genetic bottleneck. These results suggest that the invasion stemmed from a single introduction, consistent with the reported invasion history. Furthermore, the large genetic neighbourhood size and weak within-population genetic substructure suggest high connectivity across the invaded range, despite the large area occupied, and no evidence of substantial diminution of genetic diversity from the invasion core to the margins. As eradicating the species within a ~112 km radius would be necessary to prevent recolonization, eradicating Tench is likely not feasible at watershed—and possibly local—scales. Management should instead focus on reducing abundance in priority conservation areas to mitigate adverse impacts. Our study supports the argument that introduced populations can thrive despite recent bottlenecks and low levels of genetic diversity, and it suggests that landscape heterogeneity and population demographics can generate variability in spatial patterns of genetic diversity within a single range expansion.

## Introduction

Biological invasions and range expansions entail changes in population size across space and time. Following introduction to a new area, founding individuals typically carry only a fraction of the alleles present in their source population. This loss of genetic diversity, known as a genetic bottleneck, can increase inbreeding, reduce heterozygosity, and lessen the ability of introduced populations to adapt to novel environments (Chakraborty & Nei, 1977; Nei et al., 1975). Furthermore, when populations spread from restricted areas to larger regions, the number of individuals initially colonizing new habitats is likely to be limited; as a consequence, bottlenecks can occur sequentially on expanding margins (Peter & Slatkin, 2013, 2015). These serial founder events are believed to reduce the genetic diversity and the fitness of populations across space, thereby hindering their geographic spread into suitable habitat (Peischl et al. 2013; Peischl and Excoffier 2015). Yet, rather than suffering the fate of many small populations—i.e. dwindling abundance—some introduced populations expand geographically and demographically (Dlugosch & Parker, 2008; Uller & Leimu, 2011).

Significant genetic bottlenecks are expected in populations resulting from the single introduction of a relatively small number of individuals, whereas when the number of founder individuals is large, introduced populations can retain the genetic diversity of the source population (Kan & Cassel-lundhagen, 2021; Michaelides et al., 2016). Multiple introductions (spatial or temporal) are also common, yet poorly documented, and can increase genetic diversity and population structure by adding individuals and introducing new genetic variants (Dlugosch & Parker, 2008; Roman & Darling, 2007; Snyder & Stepien, 2017; Uller & Leimu, 2011). As a result of large propagule sizes and multiple introductions, the consequences of bottlenecks on genetic diversity are often modest and do not hinder the establishment of introduced species or their geographic expansion (Estoup et al., 2016; Roman & Darling, 2007).

During geographic expansions, the loss of genetic diversity from the core to the margin of the expanding range is mediated by dispersal (Ibrahim et al., 1996; Waters et al., 2013). In theory, genetic diversity losses should be exacerbated in less mobile species, owing to repetitive breeding between limited number of lineages at the expanding margin (Hallatschek & Nelson, 2008; Oskar Hallatschek et al., 2007). In addition, when long-distance dispersal events are rare, marginal populations descending from a small number of founders are likely to suffer from the genetic consequences of bottlenecks (Gandhi et al., 2016). Alternatively, highly mobile species can retain more genetic diversity because dispersal within the expanded range will contribute genetic diversity to marginal populations (Birzu et al., 2019; Goodsman et al., 2014). While often investigated using simulations and mathematical models (Andrade-Restrepo et al., 2019; Klopfstein et al., 2006; Peter & Slatkin, 2015), the outcomes of ongoing range expansion on spatial patterns of genetic diversity are not yet fully resolved empirically.

In natural populations, genetic interconnectedness is constrained by the distribution of suitable habitat and the presence of dispersal barriers. As a result, populations might expand faster in some directions than others (Samarasekera et al., 2012), and fast range expansions tend to retain higher levels of genetic diversity than slower ones (Goodsman et al., 2014). In particular, many populations exist in complex landscapes where environmental conditions (e.g. riverine flow, oceanic current) might bias the direction of dispersal and result in asymmetric gene flow (Grant et al., 2007; Lujan et al., 2020).

### A globally invasive fish, Tench, as a study system

In parts of its native Eurasian range, the Tench (*Tinca tinca*) is a cypriniform fish of conservation concern; on other continents, it is an invasive species (Avlijas et al., 2018). In eastern North America, starting from an initial importation of approximately 30 individuals sourced from Germany and stocked in a Quebec farm pond, Tench was released into the Richelieu River some time between the late 1980’s and early 1990s (Dumont et al., 2002) (Fig. 1). After an initial lag, the Tench population started growing and spreading. It was first detected in Lake Champlain in the early 2000s and subsequently dispersed towards southern Lake Champlain (southern front) and over 500 km of riverine habitat in the St Lawrence River (hereafter, the SLR) between Lake Ontario (western front) and Quebec City (north-eastern front) (Avlijas et al., 2018). Whether the Tench population resulted from a single introduction involving a handful of individuals has not been validated genetically; multiple introductions are often poorly documented, and cannot be ruled out as individuals of unknown origin have been found in another pond in the Laurentian Great Lakes watershed (Avlijas et al., 2018).

**Figure 1:**
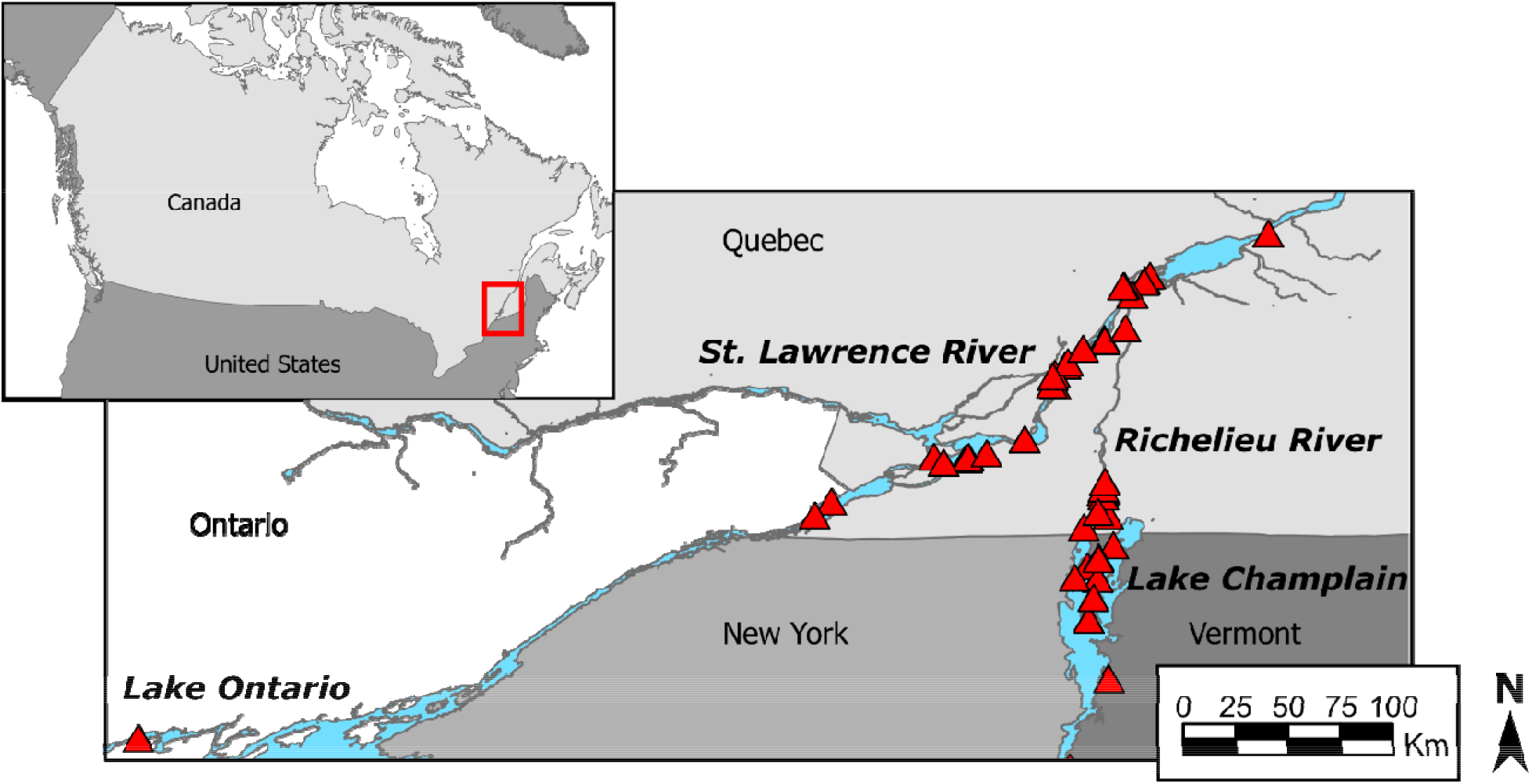
Map of the focal area in eastern North America and locations of samples (red triangles) of the contemporary population included in the genetic analysis of Tench (*Tinca tinca*).

To date, the Tench population in eastern North America has been expanding without active management by local authorities (Avlijas et al., 2018). Detailed knowledge of levels of connectivity could inform decisions regarding interventions to manage the invasion. For example, if genomic data suggests low connectivity and patches of genetically similar individuals across the invaded range, targeted culling of individuals in patches at the margins of the invasion might best limit the expansion. Conversely, if there is widespread connectivity within the invaded region, eradication might not be feasible and, thus, managers might best focus their efforts on preventing Tench dispersal to, or reducing their population density in, areas of special concern (e.g. wetlands, spawning beds) (Gandhi et al., 2016; Low et al., 2018). Finally, owing to uncertainties surrounding the levels of connectivity within the eastern North American population, the ability of Tench to disperse to new areas remains unknown.

### Study objectives

Here, we tested two contrasting hypotheses to characterize the invasion history of Tench in the region. The “bottleneck hypothesis” posits that a small number of individuals released in the Richelieu River founded the entire population. Alternatively, the “multiple-introduction hypothesis” posits that the establishment and geographic expansion of Tench resulted from more than one release event. These scenarios are characterized by contrasting levels of genetic diversity and population structure across time and space, as well as evidence, or lack thereof, for a recent bottleneck. Following conventional genomic analyses based on clustering and metrics of genetic differentiation, we computed individual- and population-based metrics of genetic diversity. To test for the occurrence of a recent bottleneck, we matched our empirical data to scenarios of bottlenecks generated using coalescent-based simulations.

We also employed powerful spatially informed population genomic approaches to test two hypotheses concerning the connectivity of the population within the invaded region. Under the “low-connectivity hypothesis”, individuals do not typically disperse over long geographic distances. If spread is constrained by landscape heterogeneity or dispersal capacity, the population might exhibit local patches of genetically similar individuals and small genetic neighbourhoods (area within which the impact of genetic drift on genetic diversity is less than that of gene flow) (Wright, 1946). Alternatively, under the “high-connectivity hypothesis”, Tench are capable of extensive movement within the established range; effective gene flow across large distances will result in a genetically cohesive population unit across the invaded range, and large genetic neighbourhood size. Finally, genetic diversity metrics should vary as a function of connectivity. Specifically, if local numbers of breeding individuals are small due to their relative isolation, which increases with geographic or ecological distances, genetic diversity metrics will vary across the invaded range.

## Materials and methods

### Data

#### Study system and sampling

Tench were sampled across the invaded range in eastern North America (Figure 1), spanning southeastern Canada (SLR, Richelieu River, and one sample from Lake Ontario) and northern Vermont, U.S. (Lake Champlain). Tench were captured using a variety of methods, including electrofishers, gillnets, fyke nets, and seine nets, from 2016 to 2019. The captured individuals were geolocated and samples for genetic analyses were collected in the form of tissue samples, preserved in 95% ethanol and held in −25°C freezers until extraction. DNA was extracted using Qiagen DNA extraction kits (Qiagen, Leusden, Netherlands). To capture spatial patterns of genetic variation across the invaded range and at finer geographic scales, we extracted DNA from a total of 345 samples collected throughout the known invaded range and quantified extracted samples with a Qubit (Thermofisher). We then sequenced those with DNA concentrations greater than 10 ng/ul (n=238). To further understand the demographic history of the population, we also extracted DNA from 40 archived fin clips collected from individuals sampled in 2002, when the species was still geographically restricted to the Richelieu River. While the DNA was degraded for most of those fin clips, we were able to include 10 samples (hereafter referred to as the original samples) in the sequencing effort.

#### Genomic data

Restriction-site-associated DNA (RAD) libraries were prepared following a three-enzyme protocol (Bayona-Vásquez et al., 2019), with modifications described in detail elsewhere (Lujan et al., 2020). Briefly, the enzymes XbaI and EcoRI were used to digest the genome and NheI to separate adapter-dimers. Isolated fragments were 340-450 bp long, ensuring that the sequencing reads (61 bp) would be at least 280 bp from all other loci, thereby helping meet the assumption of unlinked loci for downstream analyses. Libraries were sequenced on the Illumina NextSeq500 Desktop sequencer v2 for 75 bp single-end reads (Illumina Inc., San Diego, CA, USA) at the University of Toronto Center of Analysis of Genome Evolution and Function (CAGEF). We demultiplexed the resulting sequences using bcl2fastq2 v2.20 (Illumina). We then discarded low-quality reads and trimmed reads to 61 by removing heterogeneity spacers, restriction overhangs, and compensatory bases, using fastp v0.20 (Chen et al., 2018). We visualized sequence quality with FastQC v0.11.8 (Andrews, 2010) and multiQC v1.9 (Ewels et al., 2016).

Next, we filtered raw reads, assembled them *de novo*, and identified variants using the stack v2.3 pipeline (Catchen et al., 2013). For parameter optimization, we ran STACKS several times on the entire dataset, varying the values for the ustack parameter M (the number of mismatches allowed between stacks to merge them into a putative locus) from 1-5 (M1-M5) and the cstack parameter n (the number of mismatches allowed during the construction of the catalog) from M-1 to M+1. The other parameters were kept constant as they were shown to work well in previous reviews of stacks’ parameter space (Paris et al., 2017; Rochette & Catchen, 2017) and were as follows: process_radtags (--clean, --quality, --filter_illumina, -t 61, --disable_rad_check); ustacks (--disable-gapped, --model_type bounded, --bound_high 0.05, -max_locus_stacks 4, -m 3, -H); cstacks (--disable-gapped); and, population (-R 0.70, min-mac 2, --vcf). For each run, we visualized the effect of M and n values on several metrics, including the number of loci and polymorphic loci shared by 70% of the samples, the distribution of single nucleotide polymorphisms (SNPs) per loci, and the proportion of loci with proportion of heterozygotes for a given locus (H) greater than 0.55 or a read ratio deviation (D) greater than 7 inferred with HDplot (see below). Based on the effect of M and n values on these metrics (Fig S1), we identified M2 and n2 as optimal parameters as they maximized polymorphism while minimizing the number of potentially erroneous SNPs.

The resulting data were then filtered with vcftools v1.16 (Danecek et al., 2011) for data missingness. Specifically, we started with low cut-off values for missing data applied separately per individual and locus that we then iteratively and alternatively increased to exclude low-quality locus and individuals (Leary et al., 2018). Final filters included sites genotyped at >70% of individuals, individuals genotyped at >70% of loci, and polymorphic loci with a minor allele count greater than two. We imported the resulting data into HDplot (McKinney et al., 2017) to investigate allelic read depth and heterozygosity; we removed SNPs with an H greater than 0.5 or |D|>7 as they could result from potential error in loci splitting and bioinformatics. Finally, in loci with multiple SNPs, we selected the SNP with the highest minor allele frequency for downstream analyses.

### Invasion history analysis

#### Bottleneck evaluation

To test whether the Tench population experienced a genetic bottleneck, we used the approximate Bayesian Computation random-forest method implemented in DIYABC Random Forest v1.0 (Collin et al., 2021). We simulated training sets under two groups of competing scenarios referring to the absence (group 1: scenarios 1, 2, 3, and 4) or the occurrence (group 2: scenarios 5, 6, 7, and 8) of a recent bottleneck (Figure 2). Demographic parameters included four population sizes (N_ancestral_, N_bottleneck_, N_establishment_, and N_contemporary_), two sampling events t0 (for the contemporary samples) and tb (for the original samples), and up to three changes in effective population sizes (ta for changes between the contemporary and the original samples, tc for changes before the original samples, and td for changes between the ancestral and the bottleneck population). Prior values were drawn from uniform distributions with t0<ta<tb<tc<td (going backward in time). We parameterized the bottleneck timing and population size (td=[1-100]; N_bottleneck_=[2-100]) to reflect the reported importation of about 30 Tench specimens to Quebec in the 1980s (Dumont et al., 2002). Sampling events reflected the sampling of the contemporary (t0=[0-5]) and original (tb=[2-15]) population in 2017-19 and 2002, respectively. We used uniform prior values between 2 and 15 to reflect population expansion between the contemporary and the original samples (ta) and between 2 and 50 to reflect population expansion before sampling of the original population (tc). Finally, we used a broad range of priors to reflect uncertainties around population sizes (N_ancestral_, N_contemporary_, N_establishment_=[2-10,000]). Under all scenarios, we assumed that the population was the result of a single introduction (see **Results**).

**Figure 2:**
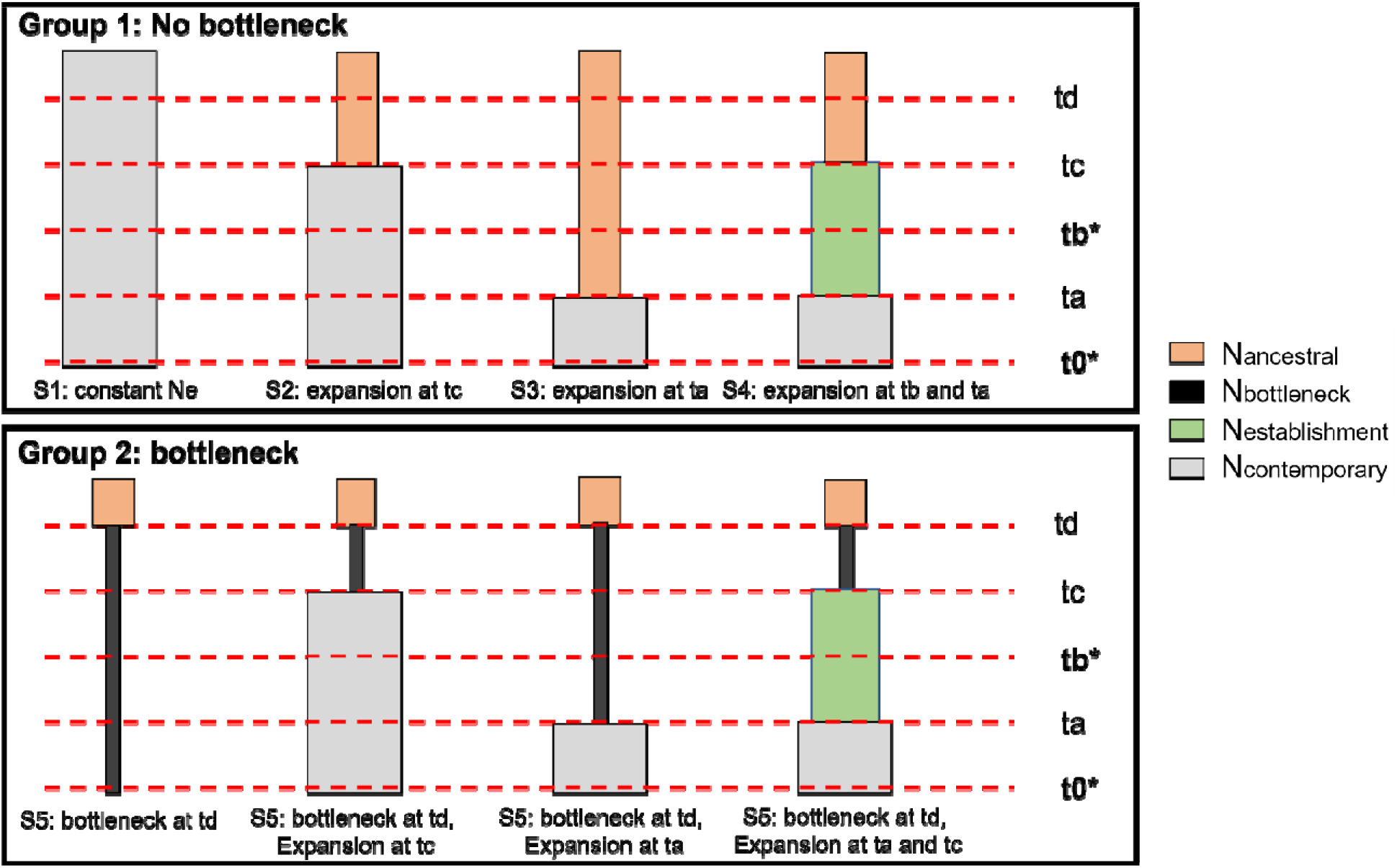
Schematic representation of the two groups of scenarios for the introduction history of Tench (*Tinca tinca*) in eastern North America tested with DIYABC Random Forest. N represents the effective population size in the ancestral, bottleneck, established, and contemporary population; t represents timing events, including sampling events (in **bold\***: t0= contemporary, tb=original population), population expansion between the contemporary and the ancestral population (ta) and between the bottleneck and the ancestral population (tc), and population bottleneck (td in group 2 of scenarios).

To identify the most likely demographic trajectory, we conducted the scenario-choice analysis twice. First, we tested whether the occurrence of a genetic bottleneck was important by conducting an analysis at the group level (with vs. without bottleneck). Second, we considered the eight scenarios separately. The training set included a total of 40,000 simulated datasets (i.e. 5,000 per scenario), and we fixed the number of trees in the constructed random forest to 1,000. Next, for scenario 5 (the best-suited scenario among the set of 8: see **Results**), we estimated the parameters involved in the invasion history: ancestral (N_ancestral_) and bottleneck population size (N_bottleneck_); and, the bottleneck time (td). For this analysis, the training set included 10,000 simulated datasets and 1,000 trees in the random forests. We ensured that the number of simulated datasets was sufficient for scenario selection and parameter estimation by evaluating the stability of both results and accuracy metrics (results not shown).

#### Temporal changes in genetic diversity and effective population size

We computed several measures of genetic diversity for both original and contemporary populations. At the population-level, we calculated observed heterozygosity (Ho), within-population gene diversity (Hs), and within-population inbreeding coefficient (Fis) using HierFstat (Meeus & Goudet, 2007). These metrics are either insensitive to (Ho, Fis) or corrected for (Hs) sample sizes. We also computed two metrics of genetic diversity at the individual-level: multilocus heterozygosity (MLH), defined as the number of heterozygous SNPs divided by the number of SNPs genotyped, was calculated using the package inbreedR (Stoffel et al., 2016); and, Individual Relatedness (IR), a metric related to the relative location of individuals along the outbred-inbred continuum, using the package Rhh (Alho & Valimaki, 2012). Negative IR values indicate outbreeding, positive values inbreeding. We tested for temporal changes between the original and contemporary populations in samples’ Ho and Hs using Wilcoxon sign-ranked test with locus treated as paired measures between populations. Because the number of samples were highly uneven between the original and contemporary populations (biased towards the contemporary population), we used randomization tests with 5,000 samples to produce accurate estimates of the p values for temporal differences in average MLH and IR.

To provide further insights into temporal changes in eco-evolutionary dynamics, we also estimated the effective population size using the linkage disequilibrium (Ne_LD_) method implemented in NeEstimator V2.1 (Do et al., 2014). This method assumes that, in small populations, heightened drift causes non-random associations between unlinked alleles. For our sample size, the least biased estimates (i.e. excluding singleton alleles) is for allele exclusion criteria Pcrit = 0.1; however, we evaluated how rare alleles affected Ne by looking at Ne variation across the range of Pcrit value (0, 0.1, 0.2, and 0.5). Stable Ne across Pcrit values are suggestive of stable, isolated populations; whereas, high variance across Pcrit values could highlight demographic processes resulting in excess of rare alleles, such as contemporary geneflow (Waples & England, 2011) or demographic expansion (Excoffier et al., 2009). We do not discuss the results from the linkage disequilibrium method in the original population as it produced infinite estimates, a likely consequence of the small sample size.

#### Spatiotemporal patterns in genetic structuring

To identify potential clusters of genetically differentiated populations across time and space, we used two nonspatial analytical approaches: a Principal Component Analysis (PCA) to explore patterns in the genetic data; and, a Discriminant Analysis of Principal Component (DAPC) using the R package Adegenet (Jombart & Ahmed, 2011) to identify clusters of genetically similar individuals. In these analyses, original and contemporary samples were included together. We evaluated the degree of population differentiation based on Fst and associated bootstrap confidence intervals. PCA creates synthetic variables to maximize variation among samples. In contrast, DAPC identifies groups of genetically similar individuals by transforming the raw data using PCA and then performing a discriminant analysis on the retained PCs to maximize between-group variability while neglecting within-group variation (Jombart et al., 2010). We used DAPC without the a-priori assumption of population structure. To assign samples to groups subsequently used as population identifier, we performed K-mean clustering from K1 to K10 with all PCs retained. To identify the best K, we used the “diffNgroup” criterion, which identifies the best K based on Bayesian Information Criterion (BIC) differences between successive values of K, as well as the “min” criterion, which retains the model with the smallest BIC. We determined how many PCs to retain based on cross validation: DAPC were performed on a training dataset comprising 90% of the samples in each subpopulation with different numbers of PCs retained and then used this to predict the group of the remaining 10%. We retained the number of PCs associated with the lower mean squared error to perform the DAPCs.

### Fine-scale population genomics and dispersal

#### Within-population subdivision

To assess contemporary population subdivision within the invaded range and identify potential areas of spatial discontinuities, we used multivariate methods that integrate spatial information into the analyses of genetic dissimilarity (i.e. Spatial Principal Component Analysis (sPCA), MEMgene). For these analyses, the original samples from 2002 were excluded. sPCA aims to identify spatial genetic patterns by analyzing spatial autocorrelation. To do so, it computes a matrix of Moran’s index inferred from the comparison between individual allelic frequencies to that of a user-defined connection network (Jombart et al., 2008). Variation is then analyzed with respect to variables (eigenfunctions) representing geographic variation that are attributed to positive (when individuals in the same neighbourhood have similar allelic frequencies, referred to as global structure) or negative (when individuals in the same neighbourhood have dissimilar allelic frequencies, referred to as local structure) autocorrelations. We set the connection network to a minimum distance neighbour graph. To characterize distance between samples, we first computed the shortest river network distance between samples that we then re-projected as Cartesian coordinates using non-metric multidimensional scaling (nMDS) in the vegan package (Oksanen et al., 2019). We then tested for significant global and local structure using the Monte Carlo simulation with 999 permutations. For visualization, we retained the two largest positive values and the three largest negative values. We performed the sPCA in Adegenet (Jombart & Ahmed, 2011).

MEMgene uses Moran’s eigenvector maps to analyse a weighted connection network. The identified spatial patterns, known as MEM eigenvectors, describe the patterns of positive and negative autocorrelation in the data (Galpern et al., 2014). The analysis then implements a forward selection procedure to identify the MEM eigenvectors that are statistically significant in a genetic distance matrix. For this reason, it performs better than sPCA in fragmented landscapes and highly mobile organisms (Galpern et al., 2014). In this analysis, we used least-cost river network distance between samples as weights in the network and the proportion of shared alleles as genetic distance. We implemented the forward selection of the statistically significant MEM eigenvectors to identify spatial patterns and used R^2^adj to estimate the strength of these spatial patterns.

#### Genetic neighbourhood size and spatial variation in genetic diversity

To understand how genetic diversity is spatially distributed across the invaded range, we used sGD (Shirk et al., 2011) to compute metrics of genetic diversity based on the genetic neighbourhood surrounding each individual. To identify the size of a genetic neighbourhood, defined as the distance at which pairwise genetic distances are no longer significantly correlated (Wright, 1946), we produce Mantel correlograms across a range of distance classes using the ecodist package (Goslee & Urban, 2007). We identified the genetic neighbourhood as the first distance class at which spatial correlation was no longer statistically significant (Shirk et al., 2011). For this analysis, we explored distance classes from 10 km to 300 km at intervals of 10 km and ran each test with 999 permutations. Next, for a set radius of 220 km (the genetic neighbourhood size based on Mantel correlograms: see **Results**), we computed observed heterozygosity (Ho), expected heterozygosity (He), and allelic richness (Ar) for each genetic neighbourhood with a minimum sample size of 20 individuals. We did not compute these metrics for the Lake Ontario sample because no other samples were within its genetic neighbourhood distance. Finally, as a comparison to the neighbourhood grouped metrics of genetic diversity, we also examined spatial variation in the individual-level metrics of genetic diversity (IR, MLH).

Next, we used linear regressions to examine the influence of range expansion on genetic diversity metrics. We included river distance (least-cost distance following the watercourse) from the putative origin, individual location relative to the putative origin (north or south), and their interaction, as explanatory variables in the models. To select the best models, we conducted backwards model selection by testing the significance of the fixed effects with likelihood-ratio tests. We started with the interaction term and removed non-significant fixed effects from the models. We removed the Lake Ontario individual whose distance from the putative origin of the invasion was 4.5-fold greater than the mean distance) and, consequently, had a disproportionate influence on the linear regressions; however, interpretation of the results were consistent across analyses including or excluding this individual. We used gdistance (Van Etten, 2017) to compute the river distance between each of the samples and the putative site of introduction in the Richelieu River.

## Results

### Genomic data

We identified 8,300 loci in the full Tench dataset; of those, 28% were polymorphic. After filtering, 1898 SNPs for 203 individuals remained in the final dataset (195 contemporary samples and 8 original samples). Many of the individuals discarded from the analysis were from dried fin clips, and the loss of those individuals is likely due to degraded DNA. The average read number per individual was 2,401,426 (SD=998,292); individual missingness was on average 11.9% (SD=7.8%) across the entire dataset, and average loci missingness was 11.9% (SD=5.9%)).

### Demographic history

The DIYABC Random Forest analysis showed clear evidence of a recent bottleneck in the SLR-Richelieu-Champlain Tench population. Classification votes and estimated posterior probabilities were supportive of scenario group 2 (1,000 votes out of the 1,000 RF-trees, posterior probability=0.996), which included the contemporary bottleneck (Table 1). When considering the eight scenarios independently, the highest classification votes and estimated posterior probabilities were for scenario 5 (820 votes out of the 1,000 RF-trees, posterior probability=0.832), which assumed a contemporary population bottleneck and no subsequent recovery. We observed that our capacity to discriminate between scenarios was better at the group-than the scenario-level, as shown by the lower overlap between the simulated datasets on the LD axes (Figure S5) and the lower global prior rate (0.029 and 0.483, respectively).

Focusing on scenario 5, parameter values had broad confidence intervals spanning most of the prior range. Estimated parameters (median [95% CI]) were as follow; 5244 [2138-9375] for the ancestral population size, 75 [37-99] for the bottleneck population size, and 52 generations [25-93] for the timing of the bottleneck event. Estimations were substantially more accurate for the bottlenecked population size and the timing of the bottleneck than the ancestral population size, as determined by lower normalized mean absolute error (NMAE) values (results not shown).

### Temporal changes in genetic diversity and effective population size

At the population-level, we detected no statistically significant difference (p=0.461) in observed heterozygosity between the original population (Ho mean (SD)=0.308 (±0.276)) and the contemporary population (Ho=0.298 (±0.196)) (Figure 3). In contrast, within-population gene diversity was significantly higher (p<0.001) in the contemporary population (Hs= 0.295 (±0.159)) relative to the original population (Hs= 0.272 (±0.200)); the change represents an average gain of 2.3% of the polymorphism between the original population and the contemporary population. We observed that Ho was higher than Hs by 3.6 and 0.3% in the original and the contemporary populations. Accordingly, Fis was negative and significantly different from zero in the original population (95% CI= [-0.172 to 0.119]) but not in the contemporary samples ([-0.020 to 0.006]). Finally, in the contemporary population, we obtained a point estimate for Ne_LD_ of 113.8 (95% CI = [106-122.5]). We observed that Ne_LD_ varied by nearly 25% across Pcrit values (FigS2).

**Figure 3:**
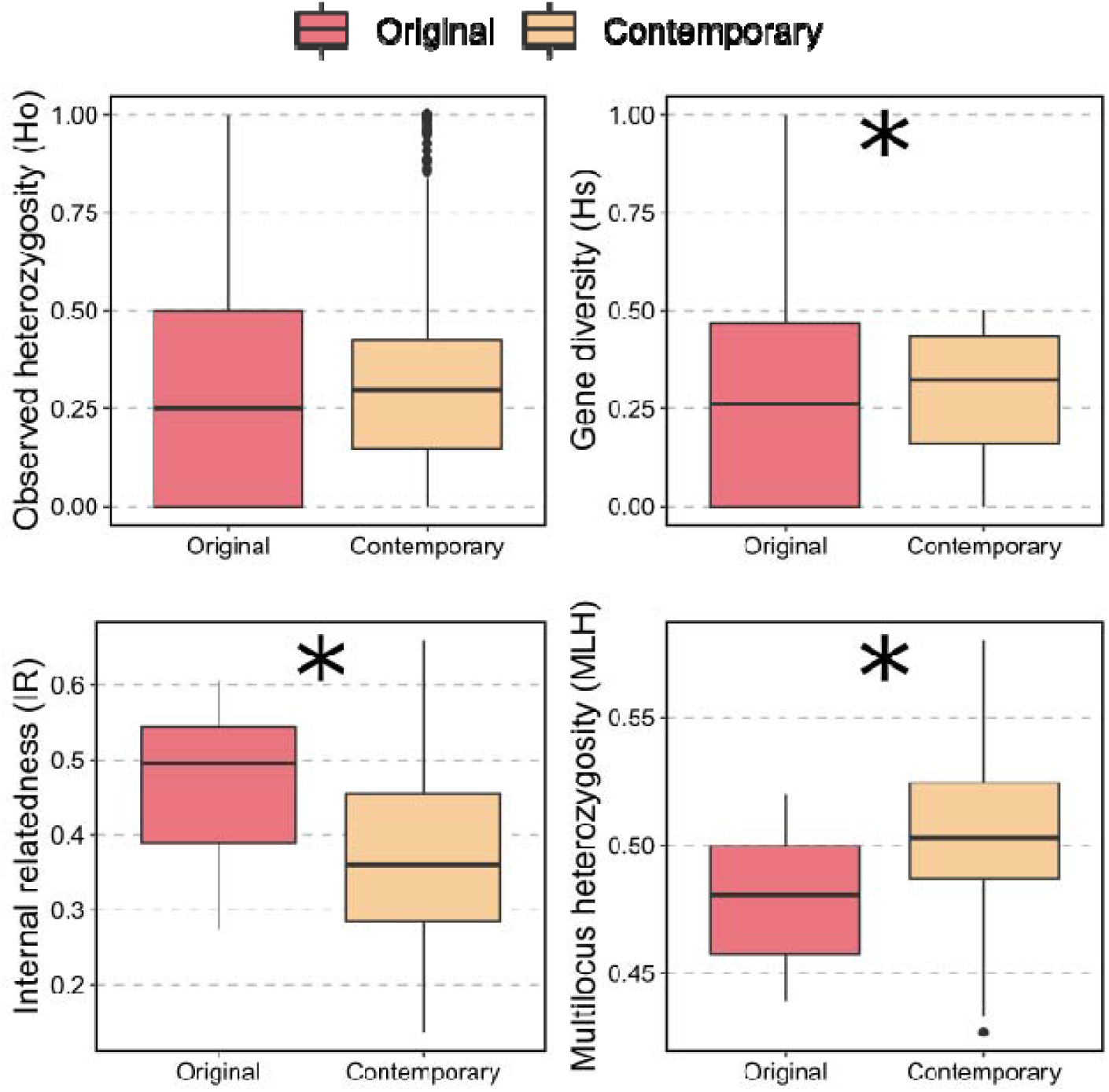
Temporal changes in genetic diversity of Tench (*Tinca tinca*) in eastern North America at the population- (Ho, Hs) and the individual-level (IR, MLH). The asterisks highlight statistically significant differences (p<0.05) in average metric values between the contemporary and the original samples.

At the individual level, genetic diversity was lower in the original population (Figure 3, Fig S3). Original population specimens had significantly (p=0.04) higher internal relatedness (mean IR (SD)=0.468 (0.109)) than the contemporary population (IR= 0.379 (0.116)) and significantly lower values (p=0.015) of multilocus heterozygosity (mean MLH_original_ (SD)= 0.479 (0.029), MLH_contemporary_ (SD)=0.504 (0.0279)). These changes represent a loss of 8.9% of the inbreeding and a gain of 2.5% of polymorphism present at the individual level between the 2002 samples and the contemporary population. The internal relatedness values were all positive, indicating that both populations show signs of inbreeding.

### Spatiotemporal patterns in genetic structuring

The PCA, DAPC, and tests of genetic differentiations indicated that the Tench population of eastern Canada, including both original and contemporary individuals, is best described as a single genetic population. In the PCA, PC1 and PC2 cumulatively accounted for less than 5% of the total variance in genetic diversity (Figure 4). While there was no clear pattern emerging from the distribution of individuals along PC2, PC1 separated Lake Champlain Tench from the rest of the samples. However, this pattern was not resolved with the de-novo DAPC analysis. Results of the K-mean clustering analysis showed that K increased linearly from 1-10 (Figure 4); while the lowest BIC score was for K=1, the greatest difference between successive BIC value was reached for K=4. However, there were no associations between the *de-novo* identified clusters and sampling locations across time and space. Concomitantly, while Fst was significantly different from zero, its value was below 1.5% (Fst [95% bootstrapped CI] =0.0067[0.0049-0.0136]).

**Figure 4:**
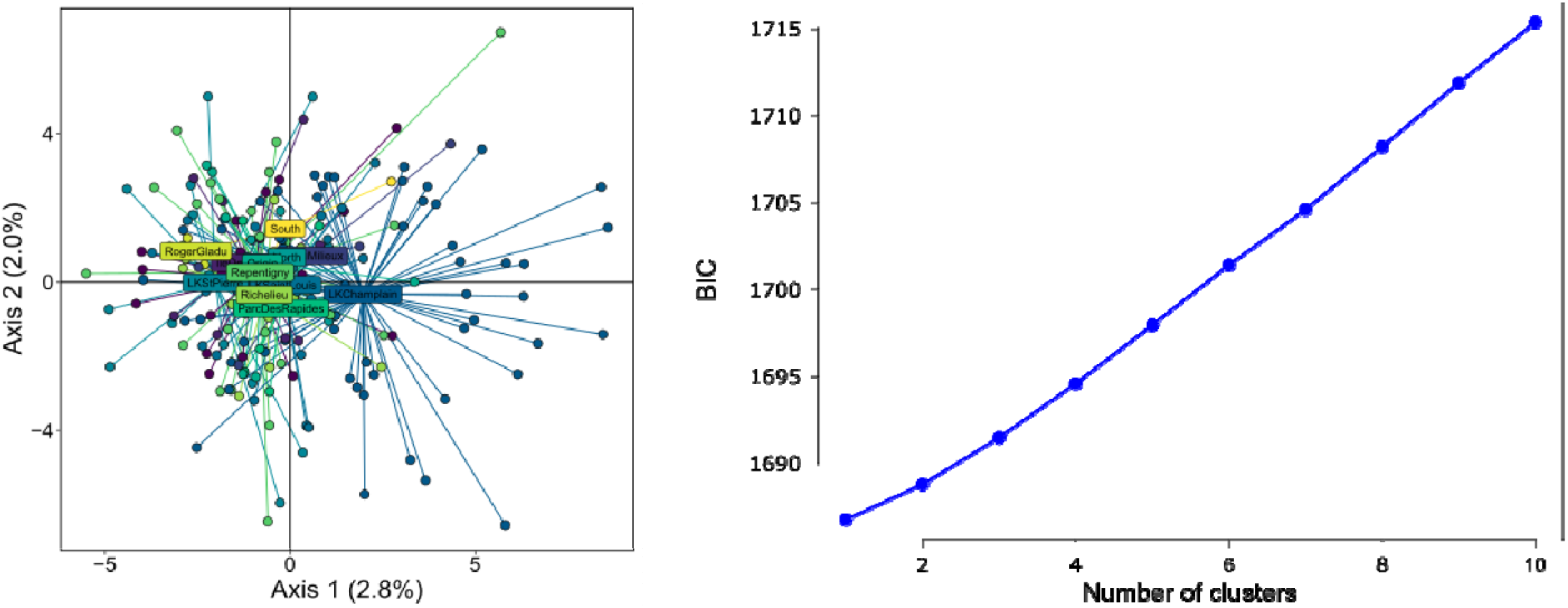
Spatiotemporal patterns in genetic structuring for Tench (*Tinca tinca*) in eastern North America was explored using several analyses. a) Principal Component Analysis showing patterns of genetic diversity distributed along PC axes 1 and 2 and the centroids of sampling locations; b) Changes in BIC values K-mean clustering to guide de-novo Discriminant Analysis of Principal Components.

### Fine-scale population genomics and dispersal

Together, the sPCA and MEMgene analysis revealed patterns of weak within-population sub-structuring across the contemporary range of invasive Tench. In the sPCA, the global test suggested the presence of statistically significant global structure across the invaded range (max(t)=0.001, p=0.001). For visualization, we retained the two largest positive values. sPCA axis 1 showed a clinal pattern of genetic differentiation across the invaded range (Figure 5). genotypes located at the front of invasion showed the most extreme scores, but there was no sharp boundaries between individuals; rather, the change was progressive, with individuals located in the middle having less extreme scores. The second axis (Figure 5) captured the same cline in allelic diversity, with a subtle difference: individuals were more similar than expected by chance in the southern section of Lake Champlain. There was no significant local structure across the invaded range (p=0.071). In the MEMgene analyses, a spatial pattern of genetic structuring emerged that was not evident in the sPCA analysis (Figure 5). Specifically, the spatial genetic pattern was more clustered, with circles of similar size and colour found in proximity, suggesting a more fragmented landscape. Overall, however, the strength of these spatial genetic patterns was weak (R^2^=0.022).

**Figure 5:**
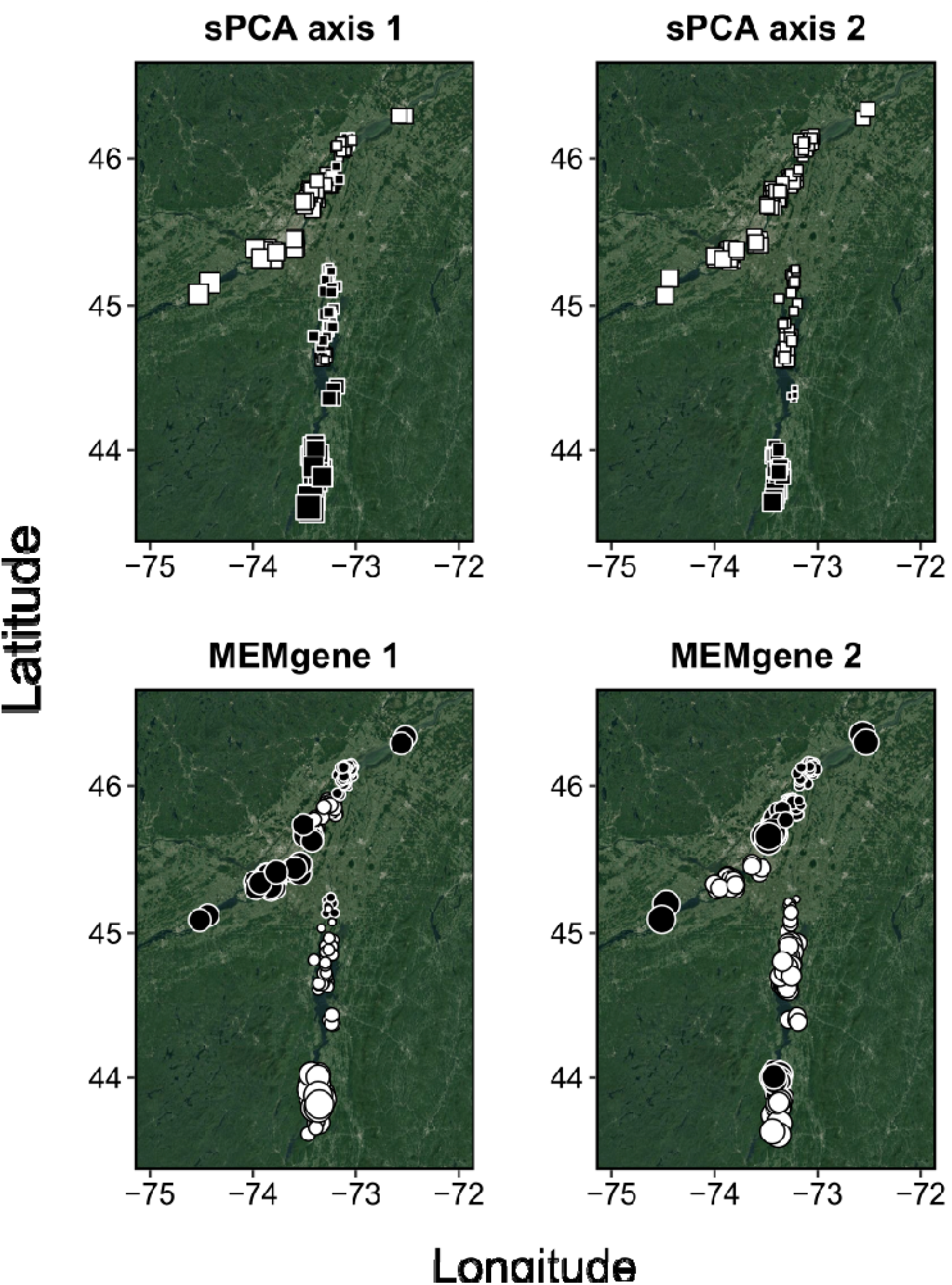
sPCA and MEMgene highlighted contrasted patterns of population sub-structuring of Tench (*Tinca tinca*) in eastern North America. Squares (sPCA) and circles (MEMgene) represent samples. To interpret the strength of genetic differentiation, square size is proportional to the eigenvalue score and colour indicates the sign: large white square are very differentiated from large black square, and small squares are less differentiated. To facilitate the visual interpretation of the plots, The Lake Ontario sample was removed.

### Genetic neighbourhood size and spatial variation in genetic diversity

The Mantel correlograms revealed the presence of significant positive spatial structure up to the distance class of 220 km (Figure 6). Estimates of genetic diversity at the genetic neighbourhood scale did not vary widely across the invaded range; mean (min, max) estimates were 1.907 (1.904, 1.910) for Ar, 0.298 (0.295, 0.302) for Ho, and −0.011 (−0.016, −0.005) for FIS.

**Figure 6:**
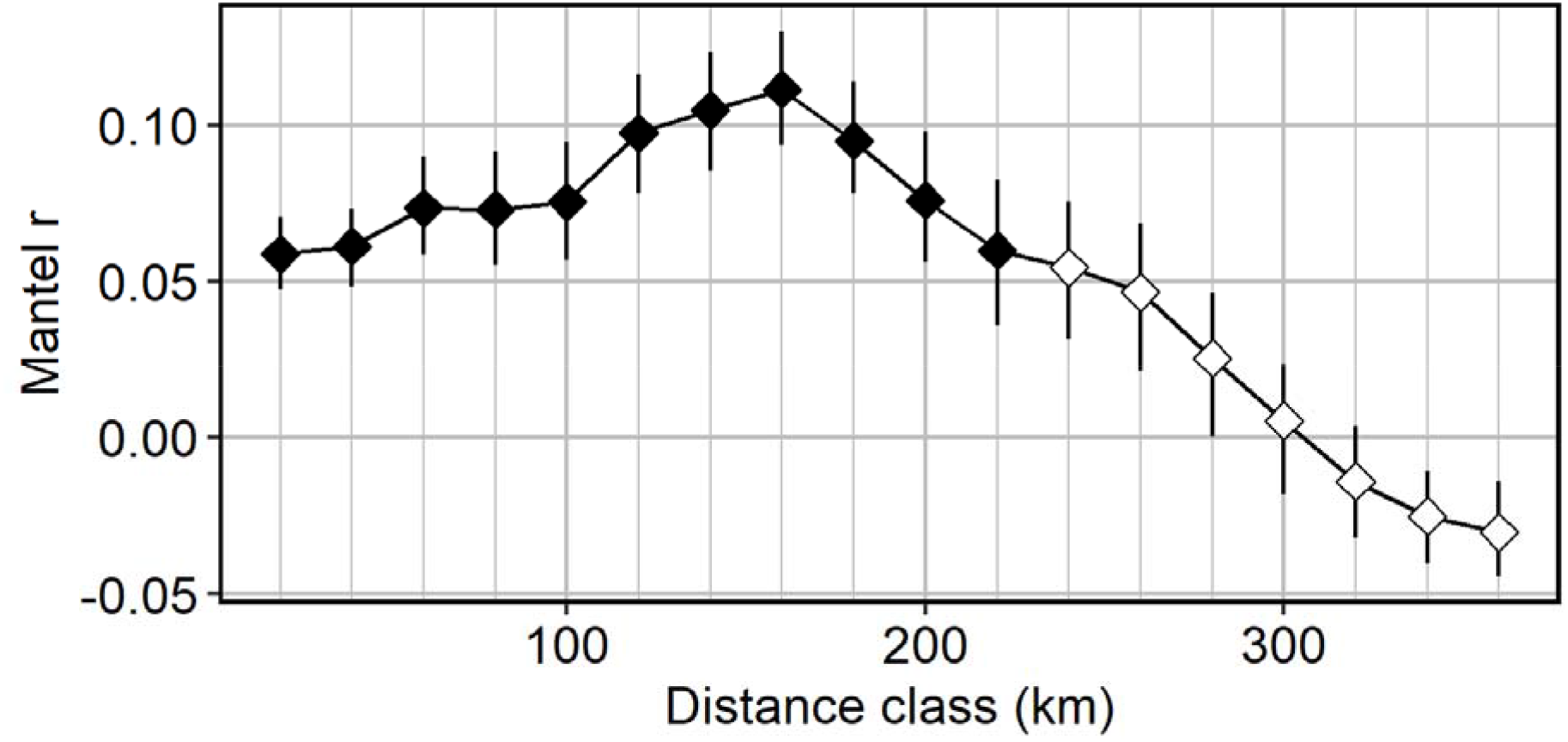
Correlogram showing spatial autocorrelation in genetic distance across a range of distance classes between Tench (*Tinca tinca*) individuals in eastern North America. The genetic neighbourhood is defined as the largest distance class with a statistically significant (indicated in black) positive correlation.

Linear models highlighted contrasting trends in genetic diversity metrics as a function of invasion directionality (whether individuals were on the southern or the northern side of the introduction site) (Figure 7). In the MLH model, the only significant fixed effect was distance (adj. R^2^(191)= 0.02, p=0.046); specifically, as distance from the origin increased, MLH decreased (−0.007 MLH/100km). In all other models, the interaction between least-cost river distance and individual location relative to the introduction site was significant. These models included the IR model (adj. R^2^(189)= 0.095, p<0.001) and, at the genetic neighbourhood-level, the Ar (adj. R^2^(189)= 0.85, p<0.001), the Ho (adj. R^2^(189)= 0.98, p<0.001), and the FIS model (adj. R^2^(189)= 0.92, p<0.001). Ar richness decreased by 0.003 and <0.001 with distance from the origin on the northern (towards the SLR) and southern (towards Lake Champlain) invasion axes, respectively. For the other metrics (ID, Ho, and FIS), trends were in opposite directions on the two invasion axes.

**Figure 7:**
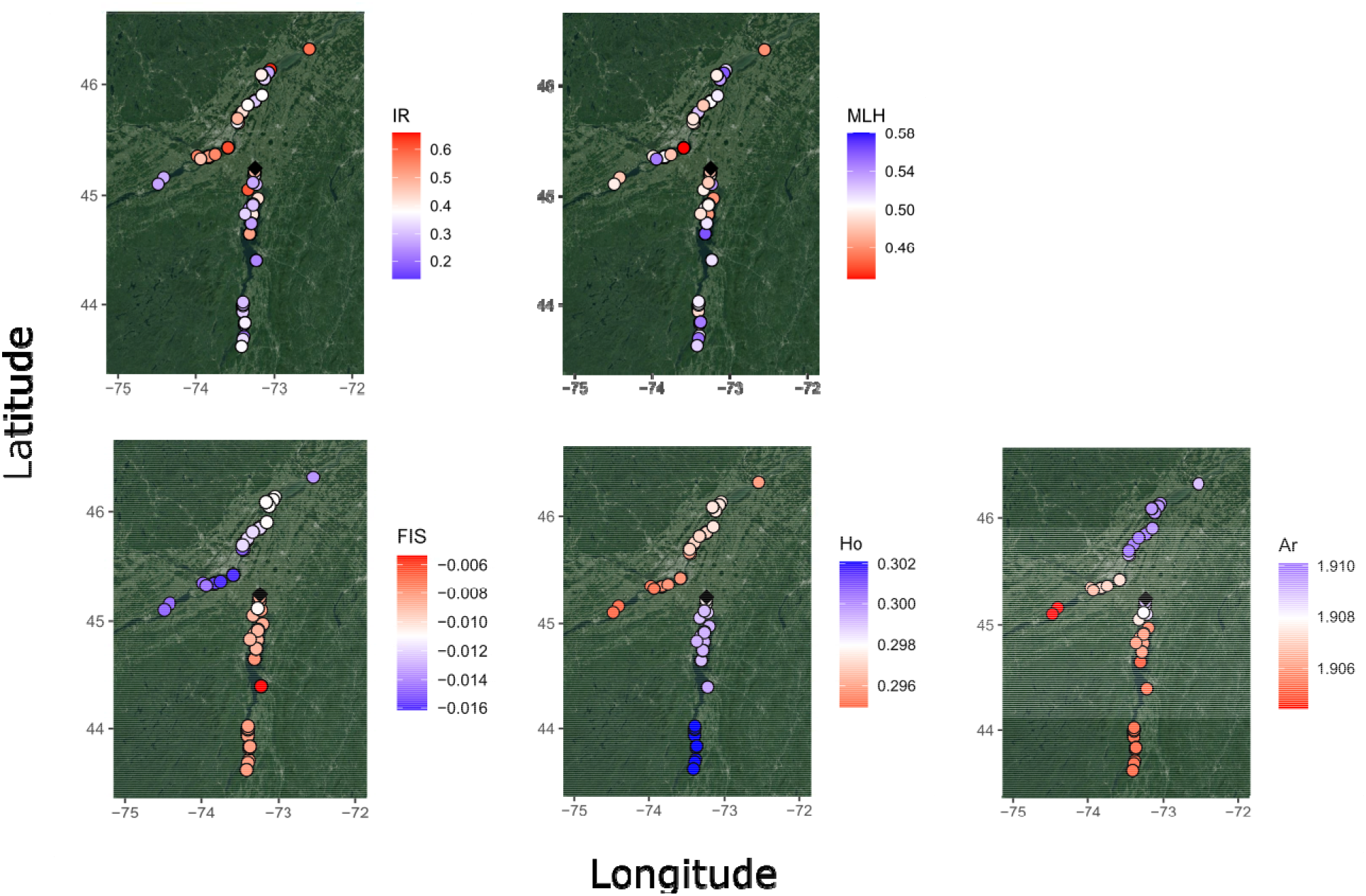
Spatial patterns of genetic diversity for the invasive population of Tench (*Tinca tinca*) in eastern North America. Genetic diversity was calculated at the individual (internal relatedness IR; multilocus heterozygosity MLH) and the genetic neighbourhood size level (inbreeding Fis; observed heterozygosity Ho; allelic richness Ar).

In general, changes in genetic diversity metrics were of small magnitude (Figure 7). Specifically, individuals were more inbred (+0.08 IR unit/100 km) as distance from the origin increased on the northern invasion axis (towards the SLR), and less inbred (−0.06 IR unit/100 km) on the southern invasion axis (towards Lake Champlain). Genetic neighbourhoods became slightly less heterozygous (−0.002 Ho/100 km) and FIS decreased (−0.004 FIS/100 km) on the northern invasion axis. On the southern axis, heterozygosity (+0.003 Ho) and FIS increased (+0.001 FIS/100 km) with distance.

## Discussion

We analyzed genomic data from the invasive population of Tench in eastern North America to discriminate between hypotheses related to the species’ demographic history and connectivity throughout the invaded range. Using a dataset with 1898 SNPs for 203 individuals, we found low genetic diversity, a lack of marked population subdivision across time and space, and evidence of a recent genetic bottleneck. Consistent with the presumed invasion history of the species in Quebec (Dumont et al., 2002; Avlijas et al., 2018), our results support the “single introduction” hypothesis and add evidence that introduced populations can thrive despite recent bottlenecks and low levels of genetic diversity (Dlugosch & Parker, 2008; Dlugosch et al., 2015). Furthermore, the weak within-population genetic substructure and extremely large genetic neighbourhood sizes exhibited by the population support the “high connectivity” hypothesis, thereby contradicting the assumption that Tench has a low capacity for natural dispersal (Moyle, 1976). Consequently, contrary to what we would expect for a species expanding its geographic range across hundreds of kilometres, Tench genetic diversity losses due to repeated founder events were not significant.

### Demographic changes and bottlenecks

We found strong evidence of a recent demographic bottleneck in the Tench population. Across eight scenarios representing several demographic trajectories with and without recent population bottlenecks, DIYABC unequivocally identified the group of scenarios incorporating a recent bottleneck as most likely. The most likely scenario included a bottleneck and no subsequent recovery, suggesting that the effective population size has not increased substantially following the recent demographic expansion. This finding was corroborated by our estimate of the effective population size in the low hundreds for the contemporary population, which likely reflects limited genetic diversity in the founder population. Furthermore, although we detected significant temporal (~6 generations) changes in some metrics of genetic diversity at the individual (e.g., MLH, IR) and the population levels (Ho), the differences were generally modest. This slow recovery in genetic diversity could be explained by the absence of immigration of new genotypes during the population recovery phase (Jangjoo et al., 2016), a hypothesis partially confirmed by the lack of strong genetic structure across time and space.

Consistent with previous studies, our results suggest that parameters inferred from DIYABC might bear some errors and downward biases, at least in point estimates (Cabrera & Palsbøll, 2017; Cammen et al., 2018). Indeed, most estimates had broad confidence intervals spanning most of the prior range. In particular, the pre-bottleneck population size was highly imprecise, with the upper bound of the range being more than 4-fold greater than its lower bound. For estimated bottlenecked population size, the lower bound of the confidence interval (37 individuals) was remarkably close to our expectations, since Dumont et al. (2002) reported that about 30 live Tench were illegally imported to QC from Germany in the mid-1980s. On the other hand, the bottleneck was estimated to have occurred as little as 25 generations ago; however, based on an age at first maturity of three years old (Ablak Gürbüz, 2011), the Tench introduction to QC occurred only ~15 generations ago. Therefore, we cannot rule out the possibility that the bottleneck occurred before the introduction of Tench (possibly at the European source) to eastern North America. Further model evaluations are warranted to ensure that the approach can be applied reliably to obtain very specific insights (e.g. population size, timing of events), as opposed to general insights (e.g. occurrence of a bottleneck), from natural populations.

Collectively, our data highlight that small, introduced populations can do well in new environments, despite bearing the genetic consequences of a bottleneck. Only thirty years after its introduction, Tench has colonized more than 500 km of riverine habitat and its population abundance has grown exponentially (Avlijas et al., 2018). Introduced populations able to flourish in novel ecosystems are often cited as examples of the genetic paradox of invasions, which suggests that they are able to adapt successfully even after experiencing genetic bottlenecks (Allendorf & Lundquist, 2003). However, additional explanations to the success of Tench in the region, despite the genetic consequences of a recent bottleneck, should be considered (Estoup et al., 2016). First, analyses based on Tench habitat requirements and life-history characteristics previously identified the invaded region as suitable for Tench (Devaney et al., 2009; Kolar & Lodge, 2002). Therefore, long-term adaptation to habitats ecologically similar to those in the native range might have facilitated the species’ successful establishment and subsequent geographic spread in the invaded region (Bossdorf et al., 2008). Second, the source of introduction of Tench to eastern North America was likely from a fish farm or a site within very close proximity of human transportation systems. Adaptation to human-altered habitats in the native range, potentially favoured by repeated introductions throughout the history of Tench (Clavero, 2019; Lajbner et al., 2011), could also be advantageous throughout the invaded region (Hufbauer et al., 2011). Indeed, the invaded area harbours two of Canada’s largest cities (Montreal, Toronto) and is characterized by highly active farming and shipping industries. Third, Tench has a relatively generalist life history and, although the SLR might not perfectly match its habitat needs, the species might have reduced needs for adaptation to become invasive (Hufbauer et al., 2011). Consequently, further research on adaptive changes is required to discover whether the population truly is paradoxical (Estoup et al., 2016). Regardless, the introduced Tench population flourishes despite harbouring low genetic diversity and having gone through a recent bottleneck.

### Range expansion and connectivity

The large genetic neighbourhood size suggests that Tench is capable of extensive dispersal across the invaded range. Tench sampled within a radius of 220 km were found to belong to the same breeding population, the first indirect genetic estimate of dispersal for the species. The geographic distance at which we detected genetic autocorrelation was consistent with the average lifetime dispersal distance of 80 km and the maximum movement distance of 250 km inferred from otolith chemistry data (Morissette et al., 2021). These convergent results of two independent studies relying on different approaches highlight the high capacity for dispersal of the species in the system.

Tench exhibited weak population substructure, most likely rooted in high dispersal ability and lack of strong landscape barriers to gene flow. Collectively, the patterns emerging from the fine-scale population substructure (sPCA and MEMgene) of Tench in eastern North America suggest a complex clinal pattern of isolation-by-distance due to limited dispersal capacity relative to the size of the landscape, with weak genetic discontinuities throughout the invaded range. The genetic discontinuities did not coincide with major known geographic barriers (e.g. dams, rapids); instead, patches tended to be linked to larger waterbodies (e.g. lakes), where individuals might aggregate and/or be sampled in higher densities.

While some species do exhibit reduced genetic diversity near the front of the range expansion compared to the core (Garroway et al., 2011; Watts et al., 2010), we found equivocal support for this theoretical prediction (Swaegers et al., 2013). Although there was a marginal loss of genetic diversity for some metrics, genetic diversity was mostly preserved during the geographic expansion of the studied population. Simulation studies predict that, during range expansions, genetic diversity losses due to serial founder events and bottlenecks can be mitigated by high dispersal from the core and/or the genetic contributions from a large number of breeders (Miller et al., 2020; Williams et al., 2019). This is likely the case in our study as the large genetic neighbourhood size and lack of strong spatial structuring are consistent with high connectivity. In particular, the large neighbourhood size indicates that dispersal throughout the entire range can occur within three generations (potentially less with occasional long-distance dispersal events). In contrast, species with limited dispersal are more likely to behave like of set of separate sub-populations, which are more likely to lose genetic diversity through genetic drift due to reduced effective population sizes (Wright, 1946). Consequently, the size of the genetic neighbourhood relative to the area of range movement, which primarily reflects species’ dispersal patterns, could be a useful predictor of the fate of genetic variation in populations undergoing range expansions.

Our results also highlight that range expansions can have different outcomes on spatial patterns of genetic diversity in populations expanding in multiple directions. We found a small, but significant, loss of genetic diversity for several metrics along the northern invasion axis (towards the SLR), while genetic diversity was preserved or increased along the southern Lake Champlain invasion axis. This result is compatible with several non-exclusive hypotheses. First, faster range expansions might retain more genetic diversity than those occurring at slower rates (Goodsman et al., 2014). Range expansion occurred faster along the southern invasion axis compared to the slower and more recent northern expansion. Second, the strength of genetic drift after colonization might influence spatial patterns of genetic diversity (Andrade-Restrepo et al., 2019; Swaegers et al., 2013). It might be that individuals in southern Lake Champlain experienced less genetic drift after colonization than those in the SLR due to earlier colonization of, and higher connectivity within, Lake Champlain, which might help retain similar levels of genetic diversity between the core populations and the southern margin. In Lake Champlain, connectivity is likely facilitated by the relatively homogeneous, nonlinear environment facilitating multi-directional dispersal. Third, differences in habitat quality might lead to local variations in population densities, thereby affecting the strength of drift and the number of mutations (Excoffier et al., 2009; Shirk et al., 2014). For example, water quality in Lake Champlain is generally higher than that of the Richelieu River due to the lower agricultural and industrial footprint and thus might be able to sustain larger populations. Collectively, this result highlights that heterogeneous landscape and population demography can generates variability in the genetic consequences of range expansions (Excoffier et al., 2009; Miller et al., 2020).

### Implications for Tench management

Our analysis of the spatiotemporal patterns of genetic diversity and structure of Tench has the potential to improve management of the invasive population in eastern North America. First, our inference of large genetic neighbourhoods (225 km) and high connectivity throughout the invaded region casts doubts on the potential for complete eradication of the species in the region. To keep areas of conservation priority free of Tench, the large neighbourhood size indicates that eradicating the species within a ~112 km radius will be necessary to prevent recolonization. Even if this could be achieved, there is still a possibility that the area might eventually be recolonized as lifetime dispersal distances up to 250 km have been documented (Morissette et al., 2021). Consequently, instead of eradication, management plans should aim at managing the species to minimize its impacts across the invaded range.

Second, our results confirm that Tench capacity for dispersal is higher than previously expected (Avlijas et al., 2018; Cudmore & Mandrak, 2011; Kolar & Lodge, 2002), which suggests that current risk assessments underestimate its potential invasiveness. This implies that colonization of the Laurentian Great Lakes is imminent (Avlijas et al., 2018; Morissette et al., 2021), as perhaps foreshadowed by the capture of a live specimen in Lake Ontario in 2018, more than a genetic neighbourhood size ahead of the known invasion front. Because this individual did not differ genetically from the rest of the invaded range, its presence in Lake Ontario might be the result of a long-distance dispersal event (natural or human-aided). Alternatively, it could indicate the presence of a sleeper population (i.e. an established population persisting in low-abundance: Spear et al., 2021) ahead of the known invasion front. Our results warrant the implementation of targeted, cohesive monitoring efforts including both conventional and eDNA sampling approaches at the invasion front near the Laurentian Great Lakes. eDNA sampling was highlighted as an efficient tool to detect Tench DNA in the area (García-Machado et al., 2021) and might be especially useful to detect the presence of a potential sleeper population, which could be below the detection threshold of conventional sampling gear (Spear et al., 2021). Targeted monitoring could enable the detection of, and rapid response to, the species when it is still at low abundance in the Great Lakes, where the species is predicted to flourish (Devaney et al., 2009).

### Study caveats

A number of issues might influence the results of our study. First, the Tench population is relatively new to the studied area in eastern North America, and it is possible that the effects of isolation-by-distance and the landscape context on population substructure require time to be realized (Anderson et al., 2010; Reding et al., 2013). However, it is worth noting that landscape structures influenced spatial genetic patterns very early (1-14 generations) in a simulation study, especially in highly vagile species (Landguth et al., 2010). Second, the Tench introduction event presumably involved a small number of founders and low genetic diversity. These characteristics could reduce our ability to detect the effects of range expansion and connectivity on spatial patterns of genetic structure (Landguth et al., 2012). Third, spatial patterns of genetic diversity and structure are simultaneously shaped by ongoing range expansion and gene flow, which are themselves influenced by landscape heterogeneity and dispersal. These processes operate on different spatial and temporal scales, and their respective influence on spatial patterns of genetic diversity are difficult to disentangle (Cushman, 2015). To confirm the results reported here, future research should employ simulation modeling implementing spatially explicit individual-based simulation frameworks, such as CDMetaPOP (Landguth et al., 2017).

## Conclusions

Understanding the consequences of founder effects and population bottlenecks for population persistence in novel environments is of great practical interest, as these eco-evolutionary challenges are commonly experienced by both invasive and endangered species (Colautti et al., 2017). We used a recently introduced invasive population as a model to examine the consequences of founder events and bottlenecks on spatiotemporal patterns of genetic diversity and structure. This study provides an example of a small, isolated vertebrate population that proliferated in a new environment despite reduced genetic diversity and a recent bottleneck (cf. Dlugosch & Parker, 2008; Uller & Leimu, 2011). Furthermore, the population did not exhibit a consistent decay in genetic diversity from the invasion core to the margins, despite the large size and habitat diversity of the invaded range. How this will impact adaptation (Excoffier et al., 2009) and dispersal (Cobben et al., 2015) as the population continues to expand into new habitats remains to be discovered. Notably, theoretical predictions suggest that, if dispersal is high enough, populations relatively well adapted to the introduced range can rapidly spread into the entire habitable range (Andrade-Restrepo et al., 2019). Range expansion itself could provide an opportunity for phenotypic changes to occur via spatial sorting, the evolution of dispersal-enhancing traits due to the concentration of fast-dispersers at the expanding front (Shine et al., 2011). However, our study also shows that, in natural settings, populations spreading in multiple directions within a single range expansion might exhibit different evolutionary trajectories. A better understanding of factors generating variability in the genetic outcomes of range expansions could allow us to make more accurate predictions related to range expansions, whether in response to introduction to a new range or to track suitable habitat conditions.

## Acknowledgements

Authors are grateful to Kunali Gohil, Francois Roy, and Emilie Simard for their help in the field; Nathalie Vachon for providing the original Richelieu samples; and, Nathan Lujan and João Pedro Fontenelle for their help with bioinformatics. This work was supported by a Vanier Canada Graduate Scholarship to T.A.B., a Natural Sciences and Engineering Research Council of Canada (NSERC) Discovery Grant (#05479) to K.M.J., an NSERC Discovery (#05226) and NSERC Strategic Project (#506528) grants to N.E.M., and a Genome Canada Large-Scale Applied Research Project grant to K.M.J. and N.E.M.

## Data Accessibility Statement

Individual genotype data and associated metadata will be made available on DataDryad upon acceptance. Code will be uploaded on github.

## Benefit-sharing statement

A research collaboration was developed between all collaborators, including scientists in academic and government agencies. Results of the research were shared with the broader scientific community. The research addresses an important topic for conservation, the rapid spread of an invasive species of major concern for native species in the St Lawrence River.

## Author contributions

S.A., J.H., O.M., and T.R. contributed samples and provided constructive feedback on the manuscript; N.M. and K.J. provided theoretical guidance and feedback on the manuscript; T.B. performed the research, analyzed the data, and wrote the manuscript.

---

**Figure S1:**
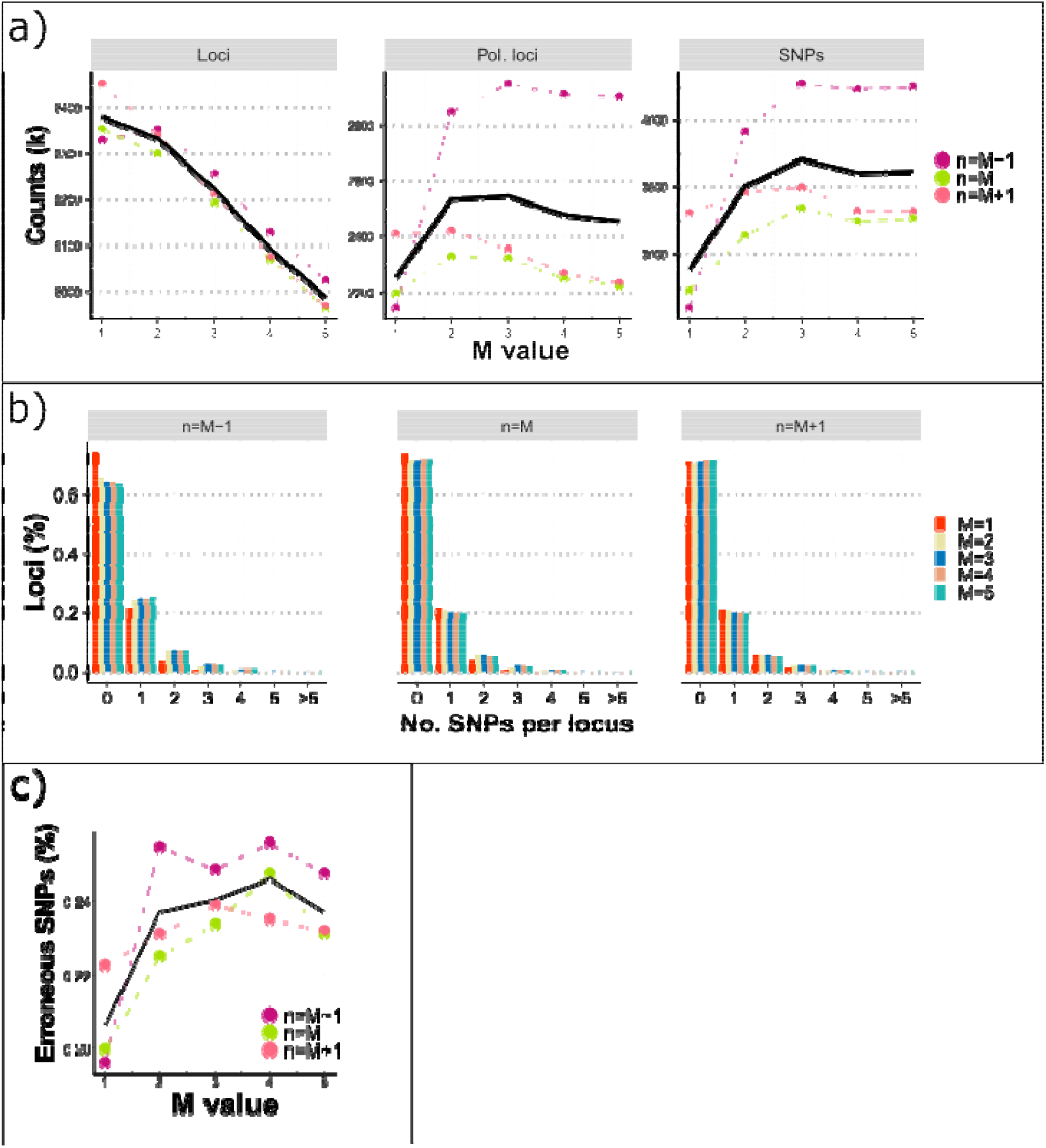
Investigation of the effect of the distance allowed between two stacks (M) and the number of mismatches in the catalog (n) in de-novo stacks assembly for Tench (*Tinca tinca*) in eastern North America on: a) the number of assembled loci present in >70% of the samples, b) the number of polymorphic loci, c) the number of SNPs, d) the number of SNPs per loci, and e) the proportion of potentially erroneous SNPs. For a given n, we observed that the number of polymorphic loci was the highest for M2 and M3 and the number of SNPs for M3. However, with M3, we observed a greater proportion of potentially erroneous SNPs and noted that there were more loci with high numbers of SNPs, suggesting that some loci might erroneously merge together for higher values of M. For a given M, the number of polymorphic loci and SNPs were greater for n=M-1; however, this parameter value was also associated with greater proportions of potentially erroneous SNPs and loci with higher numbers of SNPs. Given that we expect fixed differences to be rare in our dataset, we chose to select n=M. Accordingly, we identified M2 and n2 as the optimal parameters for our dataset.

**Fig. S2:**
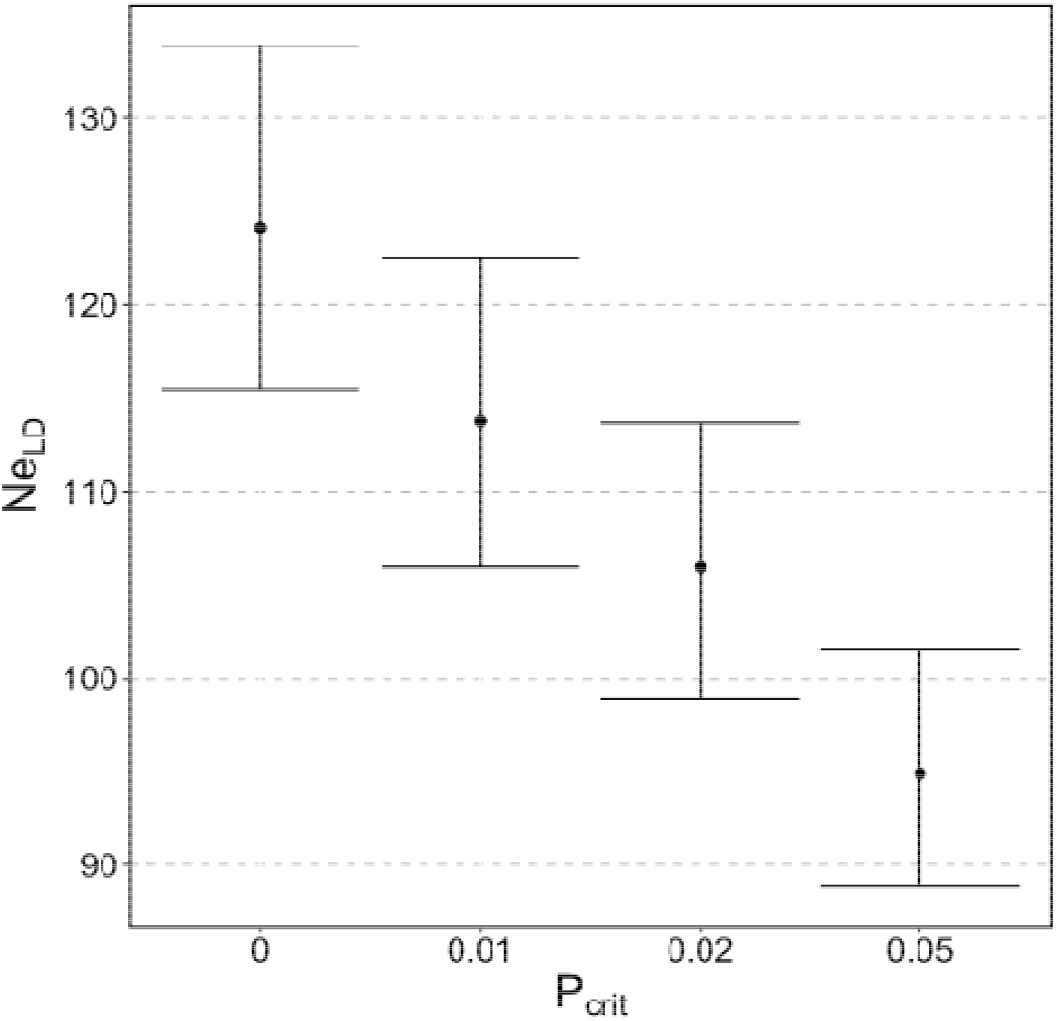
Variation in effective population size estimates from the linkage disequilibrium method (Ne_LD_) as a function of excluding rare alleles (P_crit_) for Tench (*Tinca tinca*) in eastern North America. Point estimates and associated 95% jacknife confidence intervals are shown.

**Fig S3:**
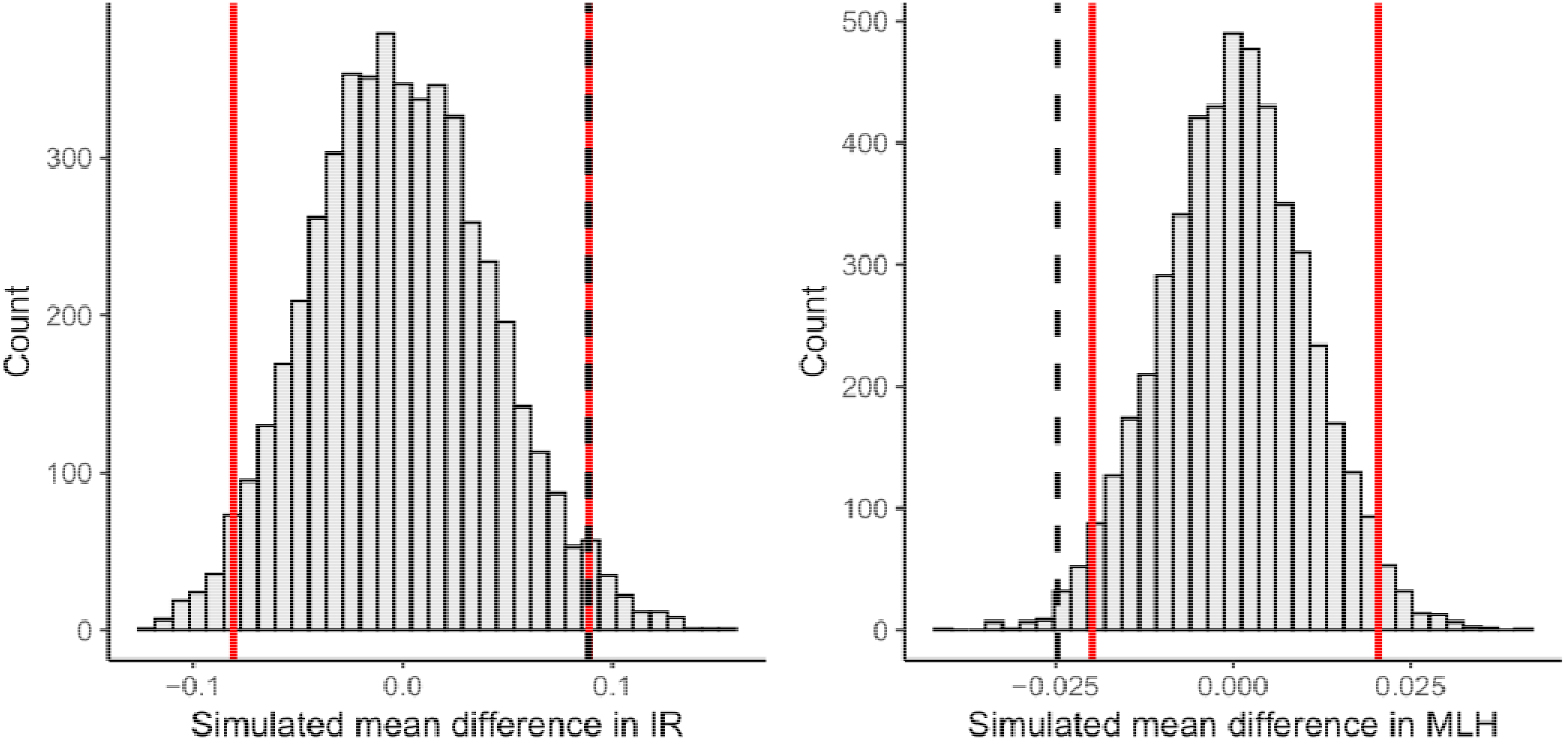
Randomization tests histogram for internal relatedness (IR) and multilocus heterozygosity (MLH) for differences between the original and contemporary invasive population of Tench in eastern North America. Red lines indicate the range within 95% of the values fall, and the dotted black line indicate the observed mean difference between the two populations.

**Fig S4:**
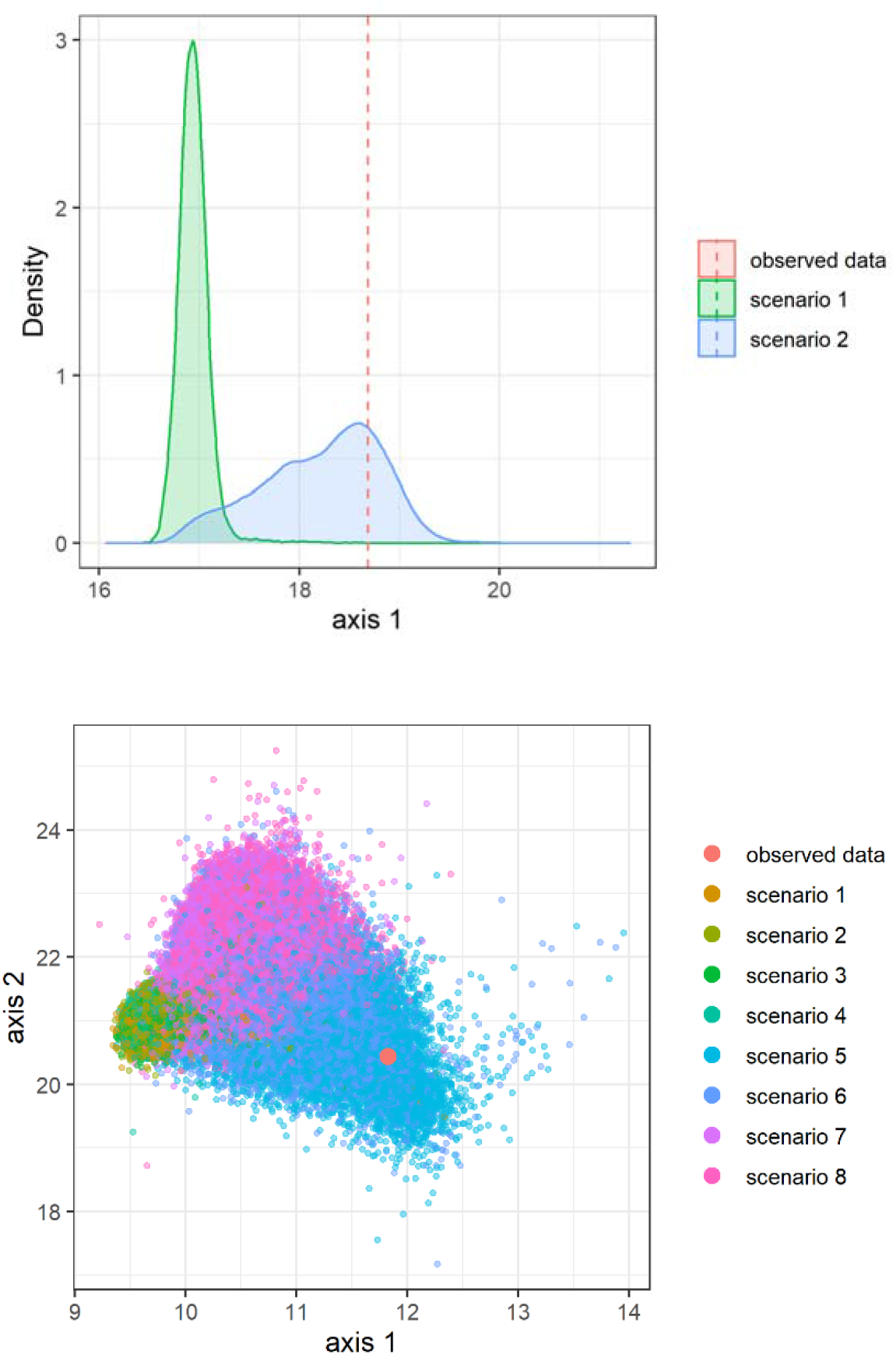
Projection of the observed Tench data on a single LDA axis for the group-level analysis (upper figure) and two LDA axes for the scenario-level analysis (lower figure). The two groups represent scenarios without (group of scenarios 1, scenario 1:4) and with bottleneck (group of scenarios 2, scenario 5:6).

